# Using a machine learning approach to complement group level statistics in experimental psychology: A case study to reveal different levels of inhibition in a modified Flanker Task

**DOI:** 10.1101/502278

**Authors:** Tanja Krumpe, Christian Scharinger, Wolfgang Rosenstiel, Peter Gerjets, Martin Spüler

**Affiliations:** Department of Computer Engineering, University of Tübingen, Sand 14, 72076 Tübingen, Germany; Leibnitz Institut für Wissensmedien, Schleichstr. 6, 72076 Tübingen, Germany

## Abstract

In this paper, we demonstrate how machine learning (ML) can be used to beneficially complement the traditional analysis of behavioral and physiological data to provide new insights into the structure of mental states, in this case, executive functions (EFs) with a focus on inhibitory control. We used a modified Flanker task with the aim to distinguish three levels of inhibitory control: no inhibition, readiness for inhibition and the actual execution of inhibitory control. A simultaneously presented n-back task was used to additionally induce demands on a second executive function. This design enabled us to investigate how the overlap of resources influences the distinction between three levels of inhibitory control. A support vector machine (SVM) based classification approach has been used on EEG data to predict the level of inhibitory control on single-subject and single-trial level. The SVM classification is a subject-specific and single-trial based approach which will be compared to standard group-level statistical approaches to reveal that both methodologies access different properties of the data. We show that considering both methods can give new insights into mental states which cannot be discovered when only using group-level statistics alone. Machine learning results indicate that three different levels of inhibitory control can be distinguished, while the group-average analysis does not give rise to this assumption. In addition, we highlight one other important benefit of the ML approach. We are able to define specific properties of the executive function inhibition by investigating the neural activation patterns that were used during the classification process.

## Introduction

In the field of experimental psychology, traditional research approaches create and test hypotheses with randomized and tightly controlled experiments. They are designed to prove or disprove that two experimental conditions are significantly different from each other. The methods of choice for this approach are group-level statistics of behavioral or psycho-physiological data usually based on averaging and analysis of variance. These methods have worked well in the past and are well established in the scientific field. However, generating detailed and descriptive models with the goal to fit these models as good as possible to a set of data, can be very problematic. These models tend to overfit to this particular set and fail to generalize to new data sets, which is one of the main reason for reproducibility issues in experimental psychology. Recently, there have been developments that try to address this problem by emphasizing the aim of prediction rather than the hypothesis-driven explanation of experimental data^1^. The reason is rather simple: Being able to predict the experimental condition from the data reliably, or even being able to predict the data based on the condition, would imply that the underlying processes can be reproduced easily.

In our opinion, both aspects, prediction as well as explanation, need to be covered in scientific approaches to understand human brain processes better. As a possible solution, we would like to suggest to establish the use of machine learning (ML) in addition to classical analysis techniques. ML could add the predictive component without losing the deep-rooted explanatory perspective^2^. In the field of machine learning over-fitting is equally an issue, but many solutions do already exist that tackle this problem. One well-known example is the implementation of a cross-validation. The data is divided into equally big parts, which, for instance, in 10-fold cross-validation, nine parts are used for training and the tenth part for testing. This step is done ten times until each part has been tested once. With this approach, it is possible to validate the generalizability of the model even with a limited amount of data. In general, most of the applications that make use of machine learning share a common goal. They focus on training models on already collected data to be able to apply the learned rules to new and unseen data.

This approach is especially interesting for applications in which the available data is densely packed with many properties or features, that can hardly be sorted and extracted manually.

An example of such a data type from experimental psychology would be electroencephalography (EEG). Standard setups usually include 32 electrodes or more from which each produces several hundreds of data points per second. It is easy to imagine, that the full spatial pattern of all electrodes contains, on the one hand, more information then can be manually explored and, on the other hand, more descriptive information for the characterization of an experimental condition than a small subset of electrodes only. Therefore, it might be a useful idea to apply ML to electrophysiological data to potentially reveal new insights, that would be missed with conventional group-level statistics on single electrodes only. Another issue that arises with EEG data and conventional group-level analysis techniques, such as averaging over all subjects, is the broad variance of brain signals between subjects. Small differences in the anatomy of brain structures can already lead to big differences in the measurable EEG signals between different subjects under the same conditions. Therefore, drawing conclusions from averages over many subjects can be misleading and not necessarily represent the ground truth regarding the underlying processes. For this problem, the ML approach equally provides a solution. It is applied subject wise on a trial by trial basis which is why inter-subject variability is not a deficit because each subject is treated individually.

All in all, we suggest that adding ML to standard analysis techniques can help to get a broader picture of the data. Aspects that have so far been neglected, such as the prediction and generalizability of the descriptive models and also the single subject level in addition to the group level of the data, can easily be integrated into the analysis to create a more comprehensive understanding of the processes. In Krumpe et al.^3^ we already successfully used this methodology to investigate the unique and shared properties of the executive functions (EFs) updating and inhibition in EEG data. To understand the significance of the results of the study, as well as the findings of the study presented in this paper, a short introduction will be given.

Executive functions are a set of processes that are essential for the control of everyday behavior. It has been established, that the EFs updating and inhibition but also shifting play a pivotal role in many recent working memory theories (e.g.,^4,5,6,7,8,9^). As a short and informal description, it can be said updating describes a process of keeping information in mind for a certain period of time^10^ and additionally being able to alter and update this information or to load new and unload older pieces of information. An everyday example would be to add and cross off items of a mental shopping list. Experimentally, updating can be tested and induced by tasks such as the n-back. The task usually consists of a sequence of items that are presented from which always the last n items constantly need to be kept in mind. Subjects need to answer for each item if it is equal or different to the item seen n-steps before. Shifting can be described as mental flexibility which relates for example to the adjustment of known rules to new circumstances^11^. Again, to give an easy example: Sorting objects once according to size and once to color, requires the adjustment of rules and certain mental flexibility to execute this task successfully. Inhibition refers to blend out irrelevant information or to avoid reflexes when the context is not appropriate^12^. One famous example to experimentally test and induce inhibition is the flanker task. In the original version a set of arrows is presented, from which only the central one should be attended. Depending on the direction the arrow displays either the right or the left button needs to be pressed as fast as possible. The remaining arrows flanking the central one can either be identical (congruent) or different (incongruent) from the central arrow and therefore, introduce a distraction that needs to be ignored. Miyake and colleagues (^13,14^) describe the relationship between the three core functions regarding unity and diversity, as they share many properties but also have distinct characteristics as individual functions. The unity and diversity of EFs as individual components of working memory has so far mainly been investigated using statistical analyses of behavioral data in either healthy subjects or patients with frontal lobe impairments (see, e.g.,^15,16,17^).

In our previous study^3^, we aimed to find out whether the shared and unshared variances of different EFs can be mapped on the one hand, on a common attentional resource in the brain or, on the other hand, on EF-specific brain functions. We chose a task design in which two EFs were induced simultaneously within the same subject, to make meaningful comparisons between the EFs. With our results, we were able to show that classification accuracies, representing an indicator for the success of the made predictions, as well as the neurophysiological interpretations of the applied ML model, lead to the conclusion that unique properties of the two EFs exist. The neural activation patterns of the model is based on a quantity of numerical values for each feature according to its importance during the classification process and therefore, according to its importance in the discriminative pattern of the experimental conditions. Hence, we have already shown once, that adding the ML component to conventional analysis techniques can reveal meaningful new insights since the classical analysis did not provide proof for that.

In the current study, we want to further focus on executive functions, but this time more on the properties of the EF inhibition specifically while emphasizing the benefits of using the proposed ML approach. We maintained the task design from the previous study in which a flanker task was combined with an n-back task. When considering the design of a Flanker task(^18,19,12^), which is well known to induce demands on the EF inhibition, it can be argued that the infrequent alternation of congruent and incongruent flanker items implies a higher level of attention and cognitive flexibility than a constant distractor item does. We, therefore, propose the hypothesis that a general readiness to execute inhibitory control is present throughout the whole task, which can be supported by theoretical explanations describing conflict adaption^20^ or selective attention^21^ processes. Based on this hypothesis, we propose that this mental state of being ready to execute inhibitory control at any time is a pre-stage of inhibition, that can be distinguished from task conditions in which this readiness is not required but also from conditions in which inhibitory control is actually executed. To evaluate this, we made small alterations to the task design already used in the previous study^3^, to be able to now distinguish three levels of inhibitory control. Since we assume that in classical flanker setups a certain amount of cognitive load is always present due to the constant readiness to execute inhibitory control, we would like to add a baseline condition in which this readiness is not required. Hence, we introduced blocks in the task design in which only congruent flanking items were presented so that no interference takes place and no inhibitory control needs to be executed. If our assumptions are correct, the distinction between the EF inhibition and true baseline demands should improve and differences between congruent trials within mixed flanker blocks (’readiness to inhibit’) compared to congruent trials in congruent only flanker blocks (no inhibition at all) can be found. Since we know that the interplay of several cognitive processes can lead to an overlap in electrophysiological signatures, or even more general, one process can overshadow the other. We aim to investigate the interplay between inhibition and updating which will be induced by an n-back task. By this, we can evaluate the stability of our hypothesis under the presence of a second executive function. The results achieved with classical group statistics and the results achieved with the ML approach will be compared to show the benefits that can be accomplished with the new methodology. Therefore, our main aim of this paper will be to promote that, using the ML approach as an additional analysis step can add meaningful information about the data that can help to interpret and understand psycho-physiological processes better.

## Results

Behavioral and neurophysiological results, as well as the performance of the ML model, will be reported in the following, to display differences between the three levels of inhibitory control. The three levels can be specified as follows: congonly describes trials from blocks in which only congruent flanker items have been presented. Trials from this category can be seen as baseline demands, in which no inhibitory control was executed. Cong describes congruent trials from blocks with mixed flankers hence, trials in which the readiness to inhibit was present but no actual inhibitory control needed to be executed, whereas incong describes incongruent trials from mixed flanker blocks, representing trials with the execution of inhibitory control. The main task of the study was to perform an n-back task on a central item, which was flanked by six items in total (three each to the left and right). The item could either be congruent (cong), which means identical to the central stimulus or incongruent (incong), which means different. Special about our task design is that an n-back was simultaneously presented with flanker items, but also the presentation of blocks in which only congruent (congonly) flanker items were presented. The two levels of the n-back task (zero and one) have been implemented to be able to evaluate the stability of the three stages of inhibitory control, during the presence of a second EF.

### Standard group-based analysis approach

#### Behavioral data

Table 1 shows the reaction time averaged over all participants and the overall accuracy of the participants responses. To display differences between the experimental conditions, the results are sorted and averaged individually for each flanker condition (congonly, cong and incong) and n-back level (zero and one). Irrespective of the task conditions, participants were able to achieve an overall task accuracy of more than 90 %. Regarding the n-back level, it can be seen that participants were consistently slower in trials from blocks with n-back level one than in blocks with n-back level zero. With one minor exception, it can also be said that the task accuracy is lower during n-back level one than for level zero. When looking at the flanker conditions, it can be seen that congonly and cong trials are answered equally fast and correct, whereas incong trials are slower and less correctly answered by the participants. Interestingly, during 1-back cong and incong answers are almost equally fast, while congonly answers have been given much faster compared to the other two categories. An ANOVA revealed that reaction time is significantly influenced by all tested factors, including participant, flanker congruency, and the n-back level. It could also be shown that there is a significant interaction between the flanker and the n-back level (see Table 2). When comparing the flanker conditions pairwise to reveal all levels of the effect, it can be seen that there are differences between the two available n-back levels (see Table 3). During n-back level zero, we find a significant difference in RT between cong and incong trials as well as between congonly and incong trials. During n-back level one we find significant differences in RT between congonly and cong trials as well as between congonly and incong trials. Interestingly no significant difference in accuracy can be found for the flanker conditions during n-back level one. During the 0-back condition, the congonly vs cong comparison is not significantly different, whereas the other two comparisons are. Even though the differences between the three levels are not consistently found across all comparisons, the analysis of behavioral data provides first clues about three different levels of inhibitory control and the interaction of the two EFs updating and inhibition.

**Table 1.** Behavioral accuracy and reaction time: Average accuracy (acc) and reaction time (RT) of the subjects categorized according to the flanker condition and the n-back level

|  |  | Flanker |  |  |
| --- | --- | --- | --- | --- |
| n-back Level |  | congonly | cong | incong |
| Zero | RT [ms] | 465.91 | 464.46 | 500.12 |
|  | Acc [%] | 94.9 | 94.6 | 90.4 |
| One | RT [ms] | 519.41 | 543.10 | 541.29 |
|  | Acc [%] | 91.9 | 91.6 | 92.7 |

**Table 2.** ANOVA on behavioral data: P-Values calculated for the average reaction times (RT) and accuracies (ACC) of all subjects per condition. An ANOVA was performed on a linear regression model, taking the participant, n-Level, flanker condition and the interaction between flanker condition and n-Level into account. Significance level has been determined to be at *p* < 0.05.

|  | Participant | Flanker | n-Level | n-Level:Flanker |
| --- | --- | --- | --- | --- |
| RT | < 0.05 | < 0.05 | < 0.05 | < 0.05 |
| Acc | < 0.05 | 0.16 | 0.15 | < 0.05 |

**Table 3.** Pairwise T-test on behavioral data: More detailed analysis of the two individual factors flanker condition and n-Level. P-Values have been calculated for the average reaction times (RT) and accuracies (ACC) of all subjects per condition with a paired t-test. Significance level has been determined to be at *p* < 0.05.

|  |  | Flanker |  |  |
| --- | --- | --- | --- | --- |
| n-back Level |  | congonly vs cong | congonly vs incong | cong vs incong |
| Zero | RT [ms] | 0.67 | < 0.05 | < 0.05 |
|  | Acc [%] | 0.70 | < 0.05 | < 0.05 |
| One | RT [ms] | < 0.05 | < 0.05 | 0.90 |
|  | Acc [%] | 0.66 | 0.28 | 0.24 |

#### Neurophysiological analysis

The grand average event-related potentials (ERPs) and spectra were calculated for the four electrode positions Fz, Cz, Pz, and O2, which were chosen as representative positions since it was shown by Scharinger and colleagues^22^ as well as in our previous study^3^, that those positions are of particular interest. Channel O2 was chosen, to cover occipital areas, since Oz was not was not recorded. While evaluating the neurophysiological signals to reveal if three levels of inhibition can be distinguished, three possible comparisons concerning were investigated (cong vs. incong, congonly vs. cong, congonly vs. incong), separately for each of the two n-back levels. The results for the ERPs can be seen in Figure 1 and the power spectra in Figure 2. Both figures show the comparisons for n-back level zero. For n-back level one the respective figures can be seen in the Appendix (Figure 5 and 6). It can be seen that at all four electrode positions almost no statistically significant differences can be found. For the power spectra, there are some indicators for differences in the congonly vs. incong comparison, but none for the other two comparisons. In consequence, the neurophysiological analysis does not support the hypothesis, that differences between different levels of inhibitory control exist.

**Figure 1.**
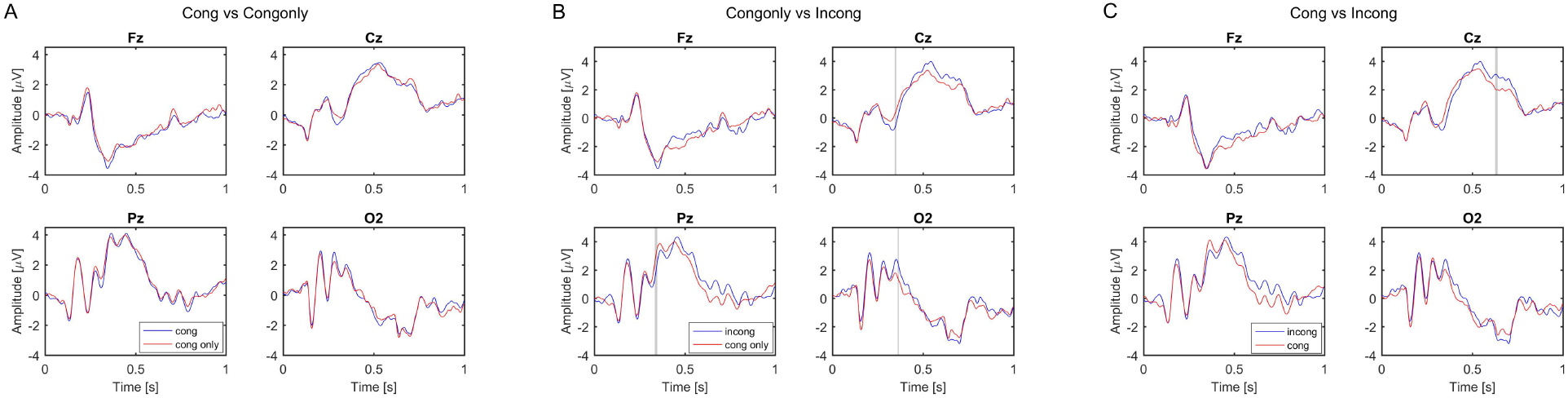
Grand average ERPs for n-back level 0: Displayed are the electrode positions Fz, Cz, Pz and O2 during n level 0. A pairwise comparison of trials with congruent, incongruent and congruent only flankers can be seen in the three subfigures. The grand average has been calculated over all 21 subjects. Grey areas indicate statistically significant differences between the two conditions (p<0.05 Bonferroni corrected, according to number of time points). A: Cong vs Congonly, B: Congonly vs Incong, C: Cong vs Incong

**Figure 2.**
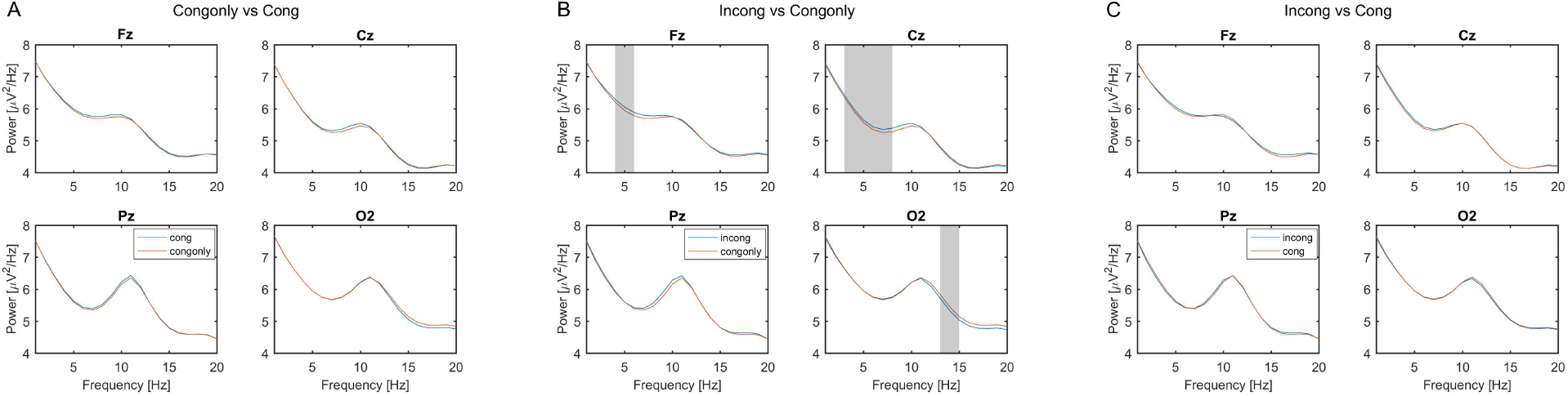
Grand average power spectra for n-back level 0: Displayed are the electrode positions Fz, Cz, Pz and O2 during n level 0. A pairwise comparison of trials with congruent, incongruent and congruent only flankers can be seen in the three subfigures. The grand average has been calculated over all 21 subjects. Grey areas indicate statistically significant differences between the two conditions (p<0.05 Bonferroni corrected, according to number of frequency bins). A: Cong vs Congonly, B: Congonly vs Incong, C: Cong vs Incong

### Machine learning approach on single subject and single trial basis

#### Classification

In the ML approach, classification is performed on single trial data to categorize each of the trials into one of the experimental conditions. The success of the classification is measured in terms of accuracy, stating how many trials out of the whole set have been categorized correctly. In the classification approach, the same comparisons as in the previous analysis steps are made. Table 4 shows the classification accuracies for the pairwise comparisons, but this time not on group-average basis but single trial and single subject basis, while taking the full spatial pattern of all electrodes into account. Therefore, trial by trial it is decided into which of the two categories it belongs. Depending on the used feature set (ERP or power spectra) the values range from 51.03 % up to 63.07 % for the distinctions. The distinction worked best for the congonly vs. incong comparison at n-back level zero and was least successful for cong vs. incong at n-back level one. The null distribution determined by permutation tests for each distinction individually revealed that most of the classification accuracies are statistically significant above chance level. The only distinction in which no statistical significance was reached with any of the used feature sets is the cong vs. incong comparison during n-back level one.

Due to this results, it can be suggested that the achieved classification accuracies provide evidence for three different levels of inhibitory control that can be distinguished.

**Table 4.** Machine Learning classification accuracy: Classification accuracies achieved with ML approach with (n-Level = 1) and without (n-Level = 0) an additional load factor. Displayed is the classification accuracy achieved with an SVM and a linear kernel during a 10 fold cross-validation. The used time frame contains 1s from stimulus onset (500 samples) from 14 channels. ERP features were additionally filtered with canonical correlation analysis (CCA), whereas power spectral features were calculated with Burgs maximum entropy method from 1–20 Hz. Statistical significance was determined by calculating an empirical null distribution with permutation tests and is indicated by *. Significance level was determined to be at *p* < 0.05

| n-back level | Features | congonly vs. cong | congonly vs. incong | cong vs. incong |
| --- | --- | --- | --- | --- |
| Zero | ERP (CCA) | 56.97 % | 63.07 % * | 57.44 % * |
|  | Power (1-20) | 59.90 % * | 60.54 % * | 51.03 % |
| One | ERP (CCA) | 54.56 % | 55.23 % | 50.11 % |
|  | Power (1-20) | 59.85 % * | 59.96 % * | 51.27 % |

#### Classification – Feature analysis

To ensure the reliability of the ML approach we use a method proposed by Haufe et al.^23^ to analyze the features that have been used in the classification approach. The method allows interpreting the features neurophysiologically, by assigning weights according to their importance within the distinction process. This way it can be controlled that only features related to the experimental condition and not caused by artifacts or features unrelated to executive control are factored in. Figure 3 shows the feature weights for the spectral features during n-back level zero. In particular for the alpha and theta frequency band which are known to correlate highly with working memory load. It can be seen that the patterns that are formed are similar in all three distinctions and only include frontal/central theta and parietal alpha components. Therefore, it is clear that the distinction is not based on noise but on features that are known for their correlation with working memory load and executive control. The ability to interpret the process of classification neurophysiologically legitimizes its use and also legitimizes the interpretation of the classification accuracies in the context of mental state characterization. Further results for n-back level one can be seen in the Appendix in Figure 7.

**Figure 3.**
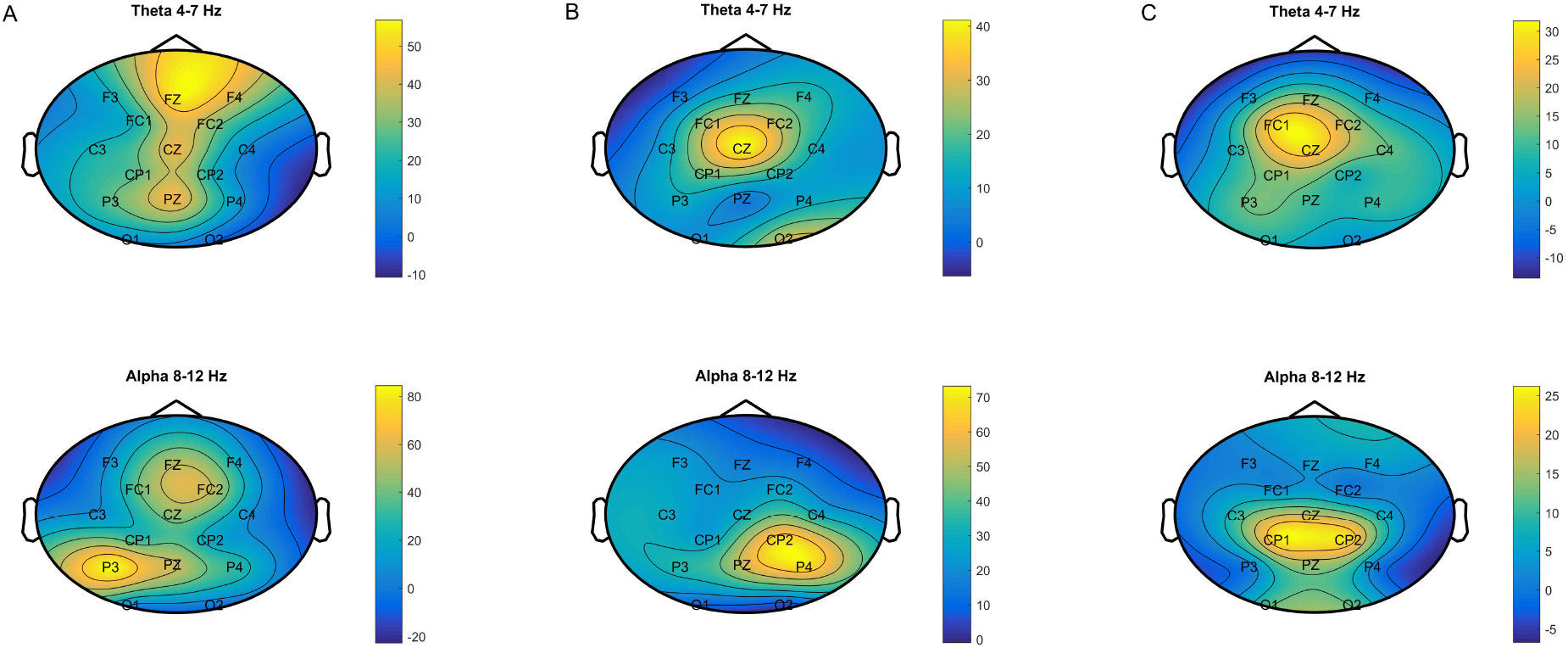
Neural activation pattern of SVM classification for n-back level 0: Displayed is the color coded activation pattern A, for the frequency bands alpha and theta in a topological distribution. The neural activation pattern has an arbitrary and undefined unit. A pairwise comparison of conflict conditions congruent (cong), incongruent (incong) and congruent only (congonly) is shown for n-back level zero. The resulting values are an average over the individual patterns of all 21 subjects. A: Congonly vs. Cong, B: Congonly vs. Incong, C: Cong vs. Incong

## Discussion

In this paper, we stated the hypothesis that three levels of inhibitory control can be distinguished in our modified version of the Flanker Task. We also stated that the addition of an ML-based prediction approach could add useful information to the classical group-based analysis approaches that are state of the art in experimental psychology. To make our point clear, we will discuss the results of the standard approach compared to the ML approach to reveal the additionally gained knowledge.

### Standard group-based analysis approach

#### Behavioral data

The general level of accuracy was high with more than 90 % in all conditions, stating that the difficulty level was adequate and all subjects were able to perform the task with sufficient accuracy. The ANOVA revealed that the n-back level and the flanker congruency have a significant interaction, as well as a significant effect on the RT. Both effects are expected according to literature^18, 24, 25^. During n-back level one, we found no differences in RT between cong and incong trials. This is an effect that can be due to a general increase of the attentional process has already been found by other authors^24^. The effect underlines the unity aspect of EFs (c.f^13^) since it can be explained by relying on a common attentional resource, based on the general activation of attentional processes that are shared. This means that based on a high attentional level no more/additional capacities can be used for the flanker processing. It can be argued that the cognitive load is lower in zero back condition and during low cognitive load enough attentional resources are available to take the flanker into account and also to solve the task sufficiently well. Hence, this could be an explanation why no difference in accuracy and RT can be found at 0-back between cong and congonly trials. It can be hypothesized that the influence of the readiness to inhibit (cong trials) on the performance seems to be minor, but gets visible when additional load is present. This would also lead to the assumption that no other confounds are introduced by using congruent flankers only if no differences between the two experimental conditions can be found during 0-back. Our approach using a block design to asses differences between congruent trials in mixed flanker blocks and blocks without flanker variation is conceptually similar to other studies^26,27^ in which the authors aimed to asses switch cost. They designed blocks in which one task was performed purely and a block in which shifting between tasks was necessary. The difference in time that was needed per block was called switch cost. A general observation that was made and can be transferred to our results is that the reaction time is slower when mixed blocks are performed in contrast to pure blocks. Theoretical validation that allows comparing congruent trials between the two blocks (congruent only vs mixed flanker block) can be found in a study by Rogers *et al*.^28^. They showed that accuracy and reaction time improve immediately after a switch trial, but no further improvement can be seen in trial three and four after the switch. Since we filter all congruent trials out that are not proceeded by a congruent trial, differences in behavioral data should not reflect an acute shifting process that influences the current congruent trial.

Using neutral flankers only would be interesting to compare with, since the do not represent competing items, but still, the use of inhibitory control might be necessary to focus solely on the central item. In addition to that using incongruent flankers only would also be interesting. This would complete the analysis regarding the question of how much influence the preparedness to inhibit, actual response inhibition and distractors without conflict have. From the behavioral results of this study, it can be concluded that two different levels of inhibitory control can be distinguished with statistical significance, but not three.

#### Neurophysiological analysis

EEG signals have been investigated on a grand-average basis at four channels of interest, covering frontal and parietal sites which are of interest concerning working memory load. No nameable significant differences have been found, suggesting that either potential differences between the experimental conditions cannot be assessed with EEG or that existing difference vanish and lose statistical significance due to averaging over 21 subjects. It could be argued that showing the results of more channels could reveal different results. However, choosing to display more channels would result in a more strict correction for multiple comparison, making it even harder to find significant differences between the experimental conditions. As already mentioned, the four chosen channels are the most important on which effects would be expected. For the averaged power spectra, the situation looks similar. Finding no significant effects there leads to the assumption that no neurophysiological differences can be found. Due to this, it needs to be stated that the classical standard analysis of ERPs and power spectra were not sufficient to draw conclusions out of the data for distinguishing three levels of inhibitory control.

### Machine Learning approach on single subject and single trial basis

#### Classification

With one minor exception classification accuracy is highest for congonly vs incong, followed by congonly vs cong trials. Accuracy is lowest for cong vs incong comparisons, independent of the n-back level. For each level distinction, statistically significant results exist, indicating that three different levels of inhibitory control can be distinguished. No inhibition, readiness to inhibit, inhibition. This fact underlines the initial assumption that, even though behavioral data does not give rise to any difference between no inhibition and readiness to inhibit at n-back level zero, statistically significant differences exist in neurophysiological data. Classification accuracy is less accurate when the n-back level is one, compared to n-back level zero, emphasizing interaction between the two conditions, which also suits the flanker effect we found relying on shared attentional resources^24^. Due to this, the non-significant performance for distinguishing cong vs incong trials during n-back level one can be explained. Therefore, it can be assumed that the higher the current cognitive load, the less accurate is the classification accuracy. Congonly trials only require the focus on one task without distraction, cong trials in mixed flanker blocks already the preparedness to inhibit potentially conflicting flankers and incong trials require the actual execution of inhibitory control. This difference can be made visible with a classification approach on EEG signals. In general, the classification approach is interesting for differentiating between different mental states or experimental conditions as it works on a single trial basis of each subject individually. Between-subject variability does not compromise the overall results as much as this is the case in standard ERP analysis. No differences in behavioral data do not mean that the experimental conditions do not induce different mental states. Essentially, it can be stated that using the classification approach in addition to the classical analysis approaches has been proven useful and generated more insights. Exploiting individual differences as a positive attribute and also the use of information from many recorded electrodes and all features that could be of relevance is one of the big advantages of the classification approach when trying to answer psychological questions.

#### Neurophysiological interpretable features

Apart from classifying the signals, it can be evaluated which neurophysiological features led to the distinction in the classification approach. This knowledge can be a useful extension to the prevailing analysis, as it gives more insight into the origin of the difference in signals between conditions. The revealed results show that only features related to working memory load have been denominated as important during the distinction. The features with the greatest importance are in the theta range, at frontal electrode positions, and in the alpha range at parietal electrode positions which are known to correlate with working memory load^29–31^. No other features have been ranked important. Therefore, it has been ensured that no artifacts or non-task related features have been factored in. For this reason, the usage of this approach can be seen as valid. Another general conclusion that can be drawn is that the same pattern, that has been found in Krumpe et al.^3^ could be replicated in this study for the EF inhibition. Despite the differences and potential high inter-subject variability, the reproducibility of the pattern reveals that this must be a rather robust pattern that has been made visible by this technique.

In our case, the usage of this approach is limited to spectral features since for temporal features a greater number of trials per condition would be required because the number of available features is a lot higher. Designing studies and acquiring EEG signals from a large number of trials would, therefore, be desirable to draw conclusions about the relevance of specific ERP features as well.

### Combination of the two methods

In direct comparison, we can see that the two methods have different strengths that cover different aspects of the data. The classical group-based statistics averages the available measures over all subjects and aims to find differences on a group level. The basis of this analysis is to find what is common between all subjects within one condition and to see if this condition is significantly different from a second condition. A priori defined hypotheses that explain the expected effects are confirmed or rejected based on the group-level statistics. The strength of this analysis is that the findings are generalized over big amounts of data that allow making inferences about the variance of the effects across subjects.

In contrast to that, the ML approach focuses on finding statistically significant differences between the two conditions for each subject individually. Instead of calculating averages or an analysis of variance, this is done by extracting mathematically optimized patterns from the available signals, that make the conditions distinguishable. To avoid over-fitting, cross-validation is implemented, that extracts the patterns from one part of the data and then validates the applicability of the found patterns on a second part of the data, that has explicitly been left out for testing. The strength of this analysis is that it allows to apply the gained knowledge to new data points. The approach predicts the condition of each data point individually which implies that the findings are generalized and validated on each subject separately that allow making inferences about the Variance of the effects within subjects.

The combination of both approaches, therefore, allows to combine an explanatory as well as a predictive approach to create more significant insights into the experimental data. Conclusions can be drawn on a group level, but also on a single subject level. Especially when dealing with EEG data, this is of great importance, as inter-subject variability can be the key to understand complex mental states on neurophysiological level. We, therefore, suggest that it is necessary to take both analysis steps into account.

## Conclusion

The here presented results could show that three levels of inhibitory control can be distinguished with the help of ML approaches in a modified Flanker task. Classical analysis approaches including the analysis of behavioral and physiological data did not create a consistent picture about the presence and differentiability of three different levels of inhibitory control. Behavioral data only gave rise to the presence of two levels, whereas the group-based averages of EEG signals were not meaningful at all. When additionally looking at the classification accuracies we found that three levels of inhibitory control can be distinguished with statistical significance. The classification approach and its validity are supported by investigating the neurophysiological interpretation of the underlying patterns that make the data distinguishable. This reveals how important it can be to also integrate single subject analysis steps, to not lose meaningful inter-subject variability. By using the results of both strategies, we combine a group-based with a single subject but also single trial analysis approach and make use of both their strengths. Therefore, a predictive component, as well as an additional explanatory component, can be gained by additionally using ML for psychological research questions.

## Methods

### Participants

21 subjects (17 females) participated in the study, for which they were reimbursed with 8 euro per hour. All subjects had normal or corrected to normal vision and no reported neurological disorders. The participants gave written, and informed consent and the study was approved by the local ethics committee. On average the subjects were 22.95 (±3.23) years old.

### Technical setup

The subjects were seated in front of a computer screen (19 inches) on which the experiment was presented by the software E-Prime (Version 2.0.10.356). A standard keyboard was used for entering the answers, by which the correctness of an answer and the reaction time were assessed. For recording EEG, a Brainproducts Acticap system with 32 electrodes was used and one BrainProducts actiChamp amplifier which was sampled at 500 Hz (PyCorder). The built-in high pass filter was set to 0.1 Hz and the low pass filter to 100 Hz. Additionally, a notch filter between 48–52 Hz was applied to eliminate power line noise. 28 electrodes were used for the recording and placed according to the extended 10–20 system^32^ (FP1, FP2, F7, F3, FZ, F4, F8, FC5, FC1, FC2, FC6, T7, C3, Cz, C4, T8, CP5, CP1, CP2, CP6, P7, P3, PZ, P4, P8, O1, O2). The ground and reference electrodes were placed on the right and left mastoid respectively with impedances below 10 kΩ.

### Task design

We used a task design of a combined presentation of flanker stimuli within an n-back task as first described by Scharinger and colleagues^22^, yet with some slight modifications. The n-back task is used to induce updating demands, whereas the flanker is used to induce inhibition demands. The experiment was presented block-wise, with six blocks in total. The main task was the n-back task with the levels 0 and 1, performed on the central item. The n-back task was performed on a set of four letters (S,H,C,F). For a schematic overview of the task see Figure 4. Yes and no answers were randomly distributed over each block with a ratio of 1:1 and given with the index finger of either the right or left hand on a standard keyboard key D and L. Which key represented the yes answer was counterbalanced throughout all subjects. Each block included 120 trials of two seconds length. One trial consisted of 500 ms stimulus presentation and a 1500 ms long blank screen, hence one block was 4 minutes long. The block order was randomized for each subject. The six blocks were subdivided into two parts between the subjects were able to take a break. By design, it was ensured that each of the two parts contains one block in which congruent flankers items were presented only.

**Figure 4.**
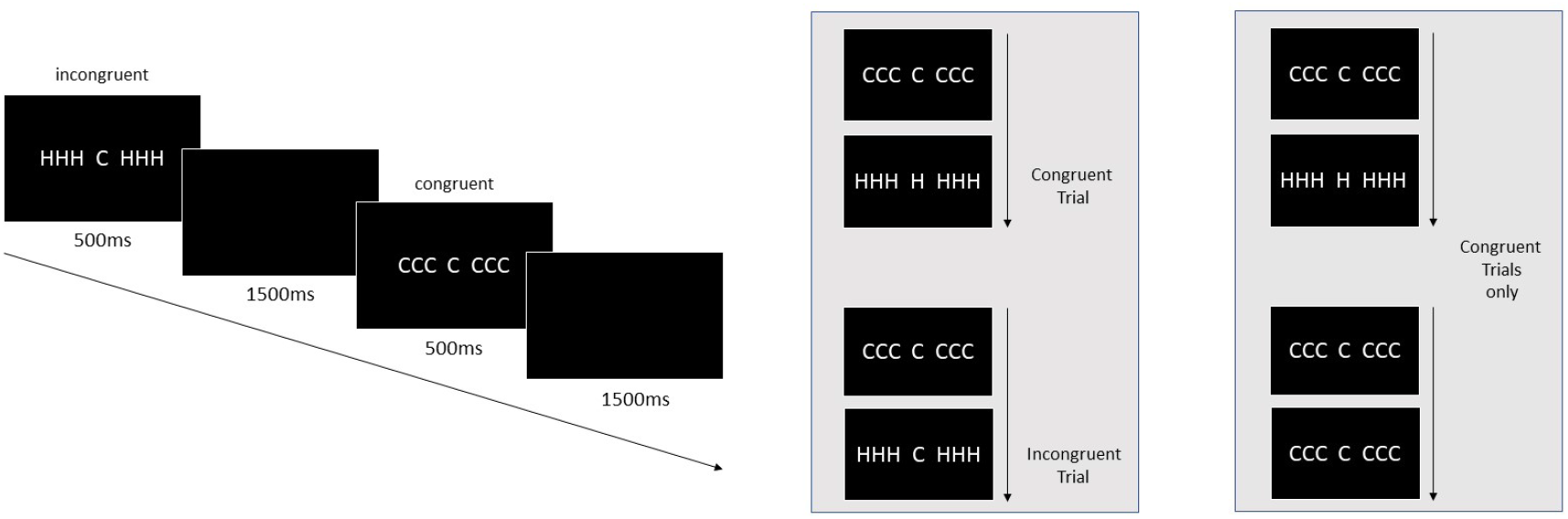
Task design: Stimuli were presented for 500 ms, followed by a blank black screen in which the subject needed to answer the n-back task on the central item with a button press. The boxes on the right show exemplary which trials were used in the analysis as congruent, incongruent or congruent only trials. The left box represents mixed flanker blocks, the right box blocks in which only congruent flanker items were presented.

The central item is flanked by six items, three to the left and three to the right. Each of the two levels was presented in three blocks of the experiment. For each n-level one block consisted of congruent flankers only (congonly), and the two other blocks consisted of alternating congruent (cong) and incongruent (incong) flankers. The congonly condition represents a baseline condition in which no inhibition is needed. The cong and incong trials are treated as separate conditions even though they are in the same block, as we presume that these trials represent two different mental processes. We propose that cong trials represent readiness to inhibit due to the unknown and unpredictable frequency of alternations of the flanker within the block, although no actual inhibition is necessary. During the incong trials, however, the flanker items need to be inhibited representing the execution of inhibitory control. Our design, therefore, enables to investigate three different levels of inhibitory control, which will be one of the main aims of this paper.

Before the experiment, each subject had to perform training to get familiar with the task. The training consisted of 2 short blocks (24 trials), one for each n-back level (0 and 1). The training blocks had to be repeated if the accuracy was below 60 % to ensure that the subject was able to solve the task correctly.

Concerning the original study, two differences need to be named which have been implemented in the current study.

### Preprocessing and analysis of the data

For the physiological data, only artifact-free trials were used for data analysis. The data was bandpass filtered between 0.5 – 40 Hz and re-referenced to common average. To remove artifacts, a threshold of ± 80 *µ*V was chosen, and all trials exceeding this level were discarded. Eye movement artifact correction was performed with a regression method by Schlögl and colleagues^33^. A pre-stimulus baseline (−100 to 0 ms) was chosen to perform a baseline correction for every trial. Stimulus onset starts with stimulus presentation in the n-back task. For the calculation of the power spectra Burgs maximum entropy method^34^ was used with a model order of 32 and a bin size of 1.

The grand average of the ERPs and spectra is investigated and to reveal statistically significant differences in the signal a Wilcoxon Ranksum Test^35^ is used to calculate a p-value. A Bonferroni correction^36^ according to the number of used tests is applied to correct for multiple comparisons. The significance level was determined to be at p < 0.05.

To be able to investigate the three levels of inhibitory control, we categorized and sorted the trials according to the presented flanker condition (congonly, cong and incong). This can be done for both n-back levels individually, leading to a total of six categories that will be investigated. Trials in the congonly condition include all correct trials from a block in which only congruent flankers were presented. Trials from the category cong, include all correct trials from blocks in which mixed flankers (alternating between congruent and incongruent) were presented. In addition to this restriction, only trials that were preceded by another congruent trial have been selected. Incong trials for that matter, include all correctly answered trials with incongruent flankers from blocks with mixed flankers, again with the restriction that only trials which are preceded by a congruent trial are used for the analysis. The categorized data will be analyzed concerning behavioral aspects like reaction times and task accuracy, as well as neurophysiologically regarding ERPs and power spectra. After assessing the task accuracy for each category and subject, only trials with correct responses have been used to calculate the average reaction times, again per category and subject. As an additional constraint, the first four trials per block were left out. Statistical significant differences between the categories have been evaluated with an ANOVA, calculated on a linear mixed effect model either on the RT or task accuracy. A pairwise t-test was performed on RT and accuracy, to reveal the specific level on which significant differences are present.

### Classification

In this paper a support vector machine (SVM)^37^ classification approach was used in addition to standard group-based analysis steps, to separate the trials according to the three levels of inhibitory control that have been defined before. A SVM with a linear kernel (C = 1)^37,38^ was applied using the LibSVM implementation for Matlab^39,40^. This approach can also be seen as a method that tries to asses the differences between two (or more) experimental conditions in EEG data. The ML approach tries to separate the data spatially within the feature space, spanned by all describable properties of the data utilizing mathematical optimization. A measure to quantify the success of the separation is classification accuracy, stating how many of all tested data points were categorized into the correct class. A pairwise separation of categories was done for each subject separately on a trial by trial basis but the overall accuracy averaged over the individual values is reported. For the classification either ERP features filtered with canonical correlation analysis (CCA)^41^ or spectral features have been used. The feature sets consist of the recordings of 14 channels (*3,*z,*4 positions) during the time frame 0 – 1000 ms starting with stimulus onset, resulting in 500 × 14 features per trial. When using the power spectra as feature sets (calculated on window 0 – 1000 ms), the frequency range from 1 – 20 Hz of the same 14 channels was used, resulting in 20 × 14 features. For each classification a 10 fold cross-validation was performed, a process in which 90 % of the data are used for training, and the remaining trials are used for the evaluation until each part of the data has been evaluated once. The average over all 10 runs is reported as the accuracy for one subject. The CCA, which is a spatial filter that can be applied to improve the signal-to-noise ratio of the EEG, is calculated on the training data and then applied on the test data in each step of the cross-validation.

Statistical testing was performed on classifications for which the accuracies were close to chance level. The statistical significance of the results was determined by permutation tests with 1000 iterations (^42,43^). The classification performance achieved in the permutations establishes an empirical null distribution on random observations, which can be used to determine significance boundaries. Therefore, in each iteration, the classification was performed in a 10-fold cross validation, but with randomly assigned class labels in the training set instead of the correct class labels. Significance level was again determined to be at p < 0.05. The original classification performance is significant when the performance values of the standard classification approach are higher than the 95 percentile of the empirical distribution calculated in the permutation test.

### Neurophysiological interpretable features

To inspect the features used for the distinction in the classification approach, a method developed by Haufe *et al*. was used that transforms the weights of the SVM classifier into neurophysiological interpretable values. This transformation is a necessary processing step since multivariate methods like SVMs combine information from several channels to improve the signal to noise ratio, thereby preventing the possibility to directly interpret the involved parameters that lead to the decision of the classifier. The step that is done in an SVM is a backward model that transforms data *x*(*n*) to the optimized and separable form *s*(*n*) by multiplying a transformation matrix on the data (Eq. 1)^23^.

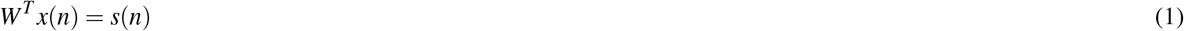

The transformation matrix represents the weights of the SVM which are mathematically optimized but cannot be interpreted regarding the neurophysiological importance of the features that are used for the distinction of classes. To reveal the individual importance, the so-called activation pattern *A* is calculated by multiplying the covariance matrices of the data (x noisy data, s separable data) with the weights of the SVM (*W* transformation matrix of the SVM, also weights), as can be seen in Eq. 2.

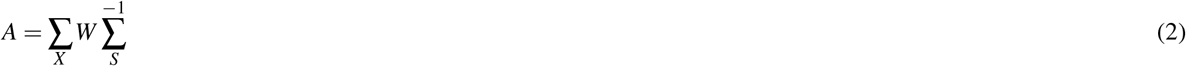

By calculating neural activation patterns, the underlying neurophysiological patterns that are responsible for the distinction can be inspected, which can provide valuable information analyzing different levels of inhibitory control.

## Acknowledgments

Tanja Krumpe is a doctoral student at the LEAD Graduate School & Research Network [GSC1028], funded by the Excellence Initiative of the German federal and state governments. The study was further supported by the German Research Council (DFG; SP 1533/2–1).

## Author contributions statement

T.K., C.S, M.S and P.G conceived the experiment, interpreted the results and reviewed the manuscript. The experiment was designed by T.K and C.S. T.K. conducted the experiment, analyzed the results and wrote the manuscript. W.R obtained funding for the research.

## Appendix

**Figure 5.**
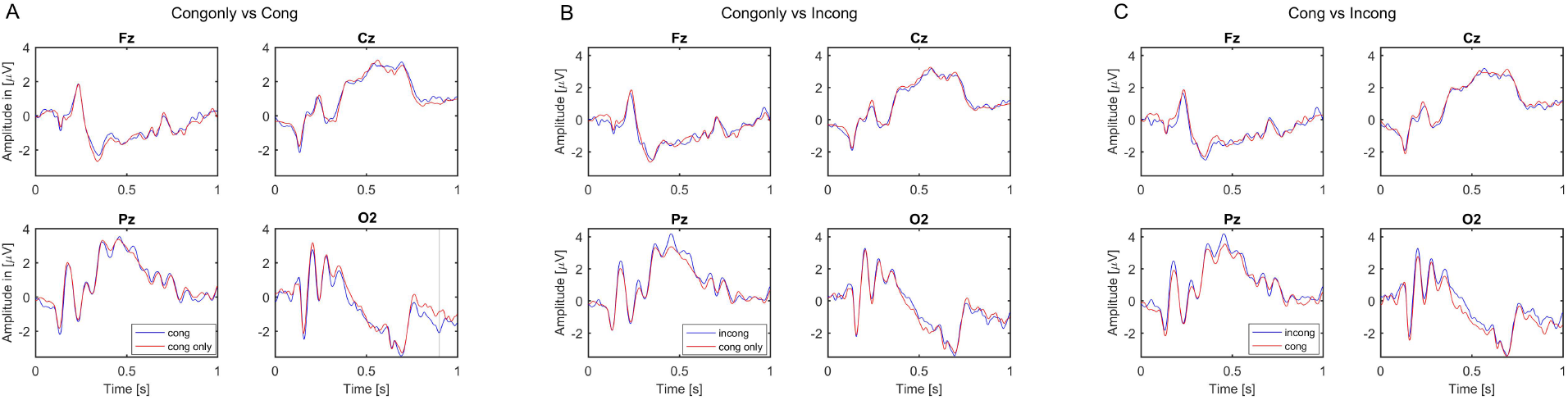
Grand average ERPs for n-back level 1: Displayed are the electrode positions Fz, Cz, Pz and O2 during n level 1. A pairwise comparison of trials with congruent, incongruent and congruent only flankers can be seen in the three subfigures. The grand average has been calculated over all 21 subjects. Gray areas indicate statistically significant differences between the two conditions (p<0.05 Bonferroni corrected, according to number of time points). A: Cong vs Congonly, B: Congonly vs Incong, C: Cong vs Incong

**Figure 6.**
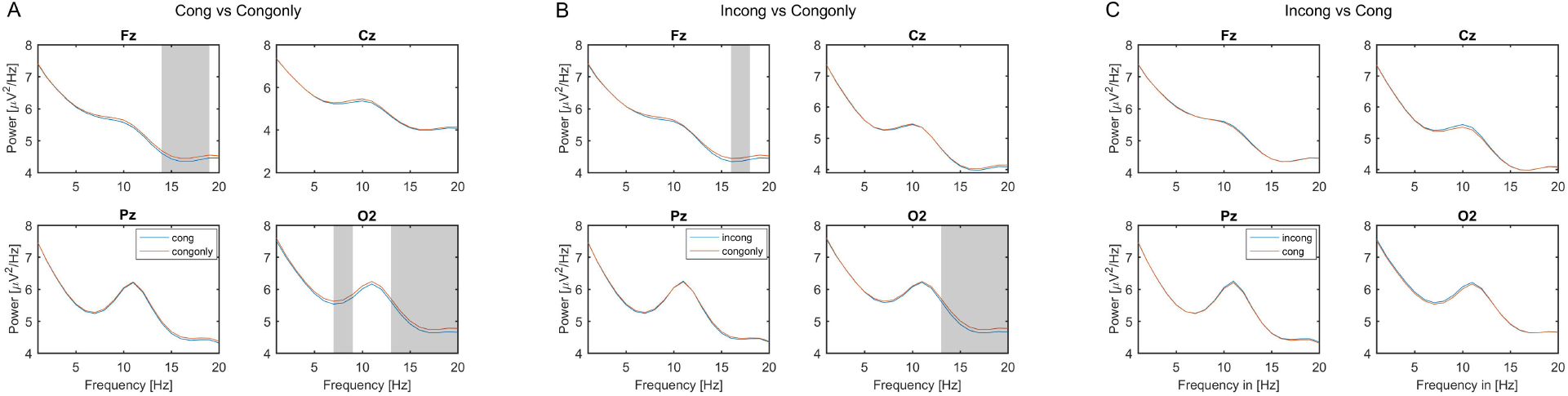
Grand average power spectra for n-back level 1: Displayed are the electrode positions Fz, Cz, Pz and O2 during n level 1. A pairwise comparison of trials with congruent, incongruent and congruent only flankers can be seen in the three subfigures. The grand average has been calculated over all 21 subjects. Grey areas indicate statistically significant differences between the two conditions (p<0.05 Bonferroni corrected, according to number of frequency bins). A: Cong vs Congonly, B: Congonly vs Incong, C: Cong vs Incong

**Figure 7.**
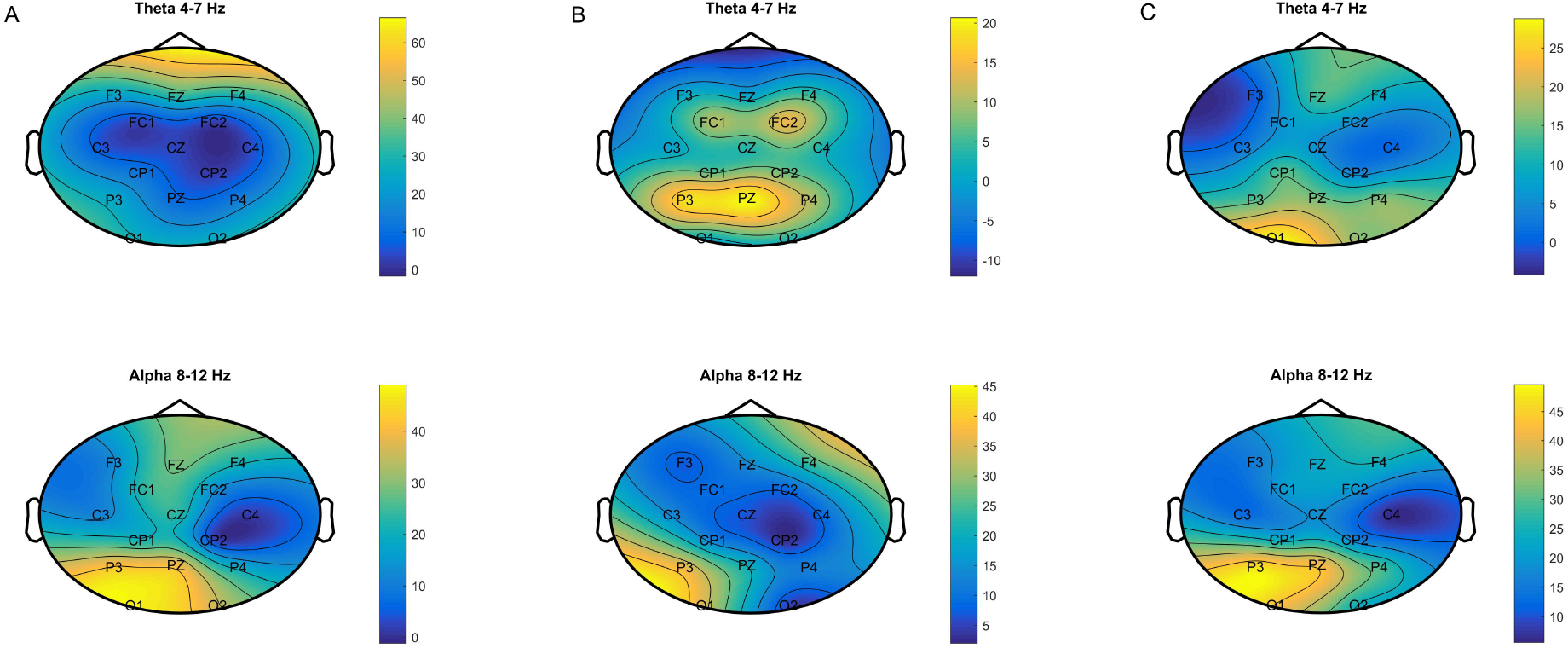
Neural activation pattern of SVM classification for n-back level 1: Displayed is the color coded activation pattern A, for the frequency bands alpha and theta in a topological distribution. The neural activation pattern has an arbitrary and undefined unit. A pairwise comparison of conflict conditions congruent (cong), incongruent (incong) and congruent only (congonly) is shown for n-back level one. The resulting values are an average over the individual patterns of all 21 subjects. A: Congonly vs Cong, B: Congonly vs Incong, C: Cong vs Incong

